# A Study of Association of ABO Blood Group types with Cancer Risk

**DOI:** 10.1101/796284

**Authors:** Vishal Singh, Upendra Yadav, Vandana Rai, Pradeep Kumar

## Abstract

More than 30 blood group systems have been recognized by International Society of Blood Transfusion (ISBT). ABO blood group is one of the most studied blood group system. ABO blood group system consist of three alleles A, B and O, out which A and B are co-dominant and O is recessive. Many researchers and investigators have found association between ABO blood group and cancer risk. It was found from the recent data that blood group A and AB is associated with increased pancreatic and gastric cancer risk. In the present study data of ABO blood group of 243 patients, both males and females, with confirmed cases of cancer was obtained from Sir Sunderlal hospital, Institute of Medical Science (IMS), Banaras Hindu University (BHU) and Apex hospital, DLW Road, Varanasi. 250 Samples of both males and females were taken as control. Out of 243 cancer patients 117 were males and 126 were females. In 243 cases enrolled in present study, highest number of cases were of breast cancer among women and lowest were rectal cancer. It was found that A blood group was associated with breast cancer, oral cancer, liver cancer and ovarian cancer as compared to other blood group and blood group O was associated with lung cancer, gastric cancer, colon cancer, skin cancer and endometrial cancer.

## Introduction

ABO blood group system contains three antigens (i.e. A, B and H) and is clinically most important blood group system among 33 blood group systems (1). Blood groups classification refers to the antigens present or absent on the red blood cells (RBCs) surface. The gene for ABO is located on chromosome 9 at 9p34.1-q34.2. ABO gene has 7 exons. ABO locus has three main allelic forms A, B and O. The frequency of A and B blood groups differs among the population of the world (2, 3, 4). Several studies have been carried out to find the frequency and association of ABO blood groups with different types of diseases in different population of the world (5–34)

**Table.1:** Association of ABO Blood Group with Different Type of Cancer-

| **S.N.** | **Cancer** | **Sample Size** | **Blood Group Association** | **Country and State** | **References/ study** |
| --- | --- | --- | --- | --- | --- |
| 1 | Breast Cancer | 1713 | A | Korea | Park et. al., 2017 |
| 2 | Breast Cancer | 206 | A | Rajasthan | Saxena et. al., 2015 |
| 3 | Breast Cancer | 166 | A | Greece | Meo et. al., 2017 |
| 4 | Breast Cancer | 197 | A | Iran | Shiryazadi et. al., 2015 |
| 5 | Pancreatic Cancer | 166 | A | Germany | Pelzer et. al., 2013 |
| 6 | Pancreatic Cancer | 633 | A | Turkey | Engin et. al., 2012 |
| 7 | Pancreatic Cancer | 627 | A | Germany | Rahbari et. al., 2012 |
| 8 | Pancreatic Cancer | 274 | A | U.S | Greer et. al., 2010 |
| 9 | Liver Cancer | 88 | A | Bangladesh | Hosen et. al, 2018 |
| 10 | Gastric Cancer | 1412 | A | China | Xu et. al., 2016 |
| 11 | Gastric Cancer | 1045 | A | China | Wang et. al., 2012 |
| 12 | Gastric Cancer | 3245 | A | Korea | Song et. al., 2013 |
| 13 | Colorectal Cancer | 1620 | A | Turkey | Urun et. al., 2012 |
| 14 | Esophageal Cancer | 480 | A | India | Kumar et al., 2014 |
| 15 | Lung Cancer | 307 | A | Turkey | Oguz et. al., 2013 |
| 16 | Lung Cancer | 2044 | A | Turkey | Urun et. al., 2013 |
| 17 | Lung Cancer | 458 | A | Jordan | Alqudah et al., 2018 |

## Data Collection

Data was collected from Sir Sunderlal Hospital, IMS, BHU, Varanasi and Apex Hospital, Varanasi. Out of 243 confirmed diagnosed cancer patients, 117 were males and 126 were females. Out of which 57 sample were patients suffering from breast cancer, 17 were patients suffering from kidney cancer, 29 were patients suffering from lung cancer, 27 were patients suffering from oral cancer, 9 were patients suffering from liver cancer, 21 were patients suffering from blood cancer, 6 were patients suffering from brain tumor, 3 were patients suffering from prostate cancer, 10 were patients suffering from endometrial cancer, 17 were patients suffering from ovarian cancer, 18 were patients suffering from gastric and stomach cancer, 17 were patients suffering from colorectal and colon cancer, 2 were patients suffering from Rectal and anal canal node and 10 were patients suffering from skin cancer. Data of 250 Samples were taken as control. Out of which 160 were male and 90 were female.

## Results

Blood group A was highest among the total 243 cancer patients. The distribution of ABO blood groups among the patients suffering with breast cancer were blood group type A(49.12 %), blood group type B(15.78 %), blood group type AB (8.77 %) and blood group type O (26.31%). Blood group A was found more in patients suffering from breast cancer as compared to the other blood groups (P value 0.0005).

**Table.2:**
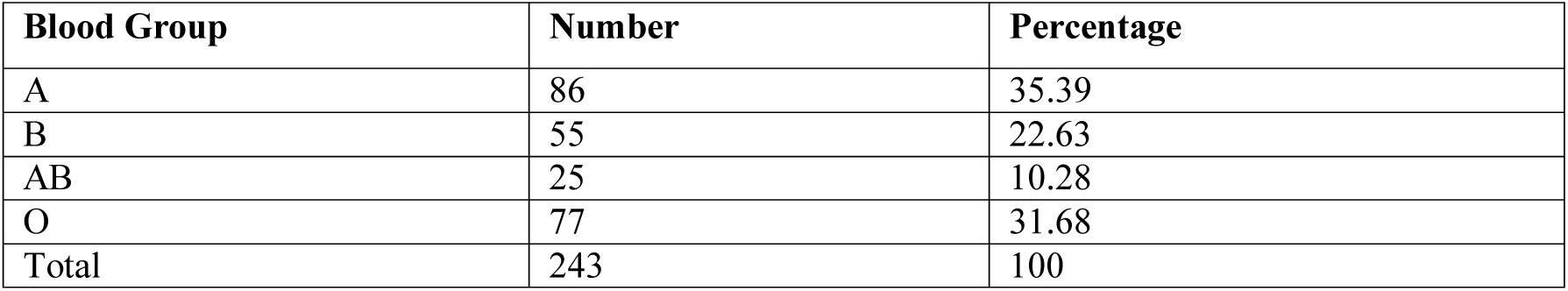
Distribution of ABO Blood Group in Cancer Patients (n=243).

| Blood Group | Number | Percentage |
| --- | --- | --- |
| A | 86 | 35.39 |
| B | 55 | 22.63 |
| AB | 25 | 10.28 |
| O | 77 | 31.68 |
| Total | 243 | 100 |

**Figure 1.**
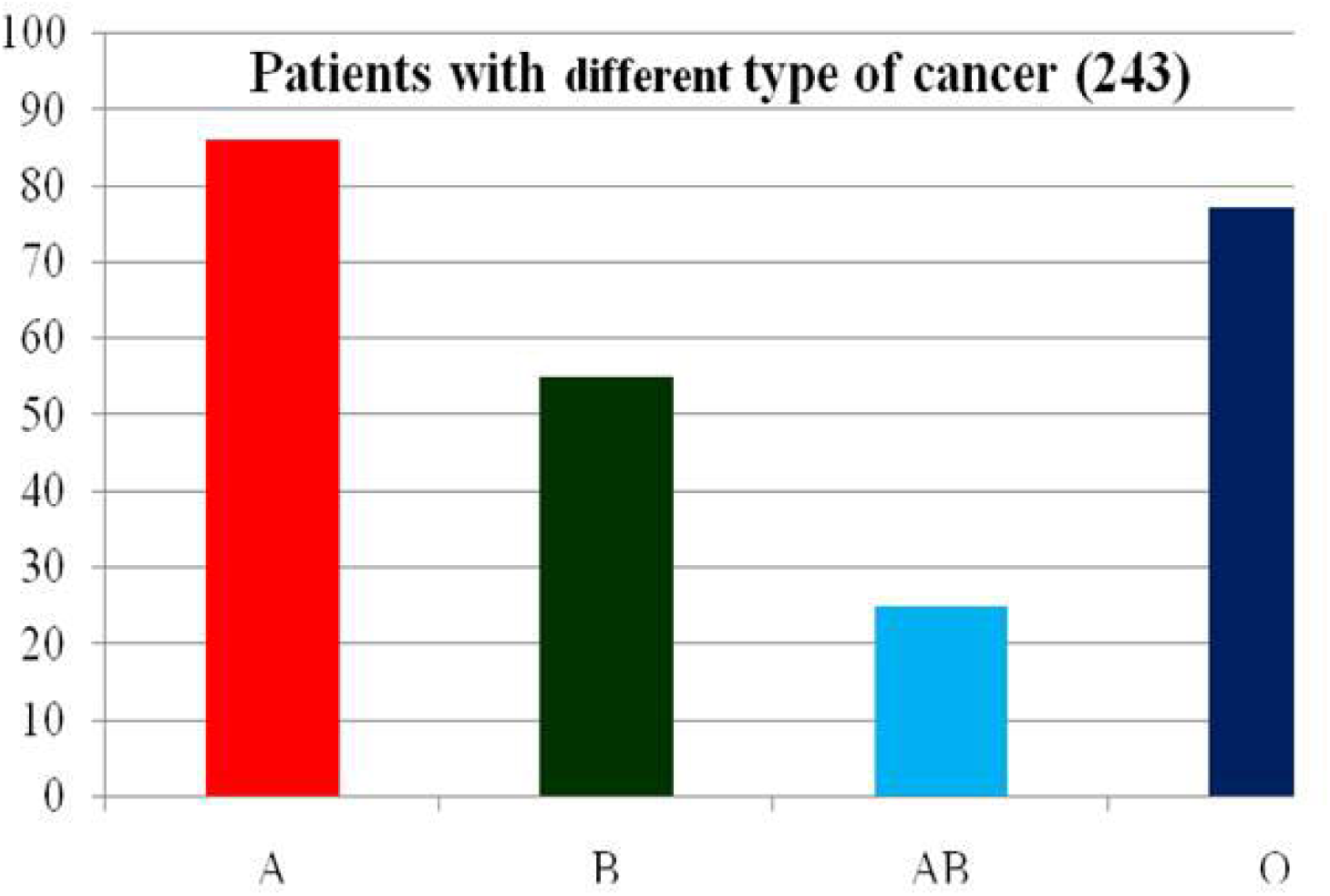
Distribution of ABO Blood Group in Cancer Patients(n=243).

**Table 3.** Distribution of ABO blood group in normal population.

| Blood Group | Number | Percentage |
| --- | --- | --- |
| A | 45 | 18 |
| B | 94 | 37.6 |
| AB | 25 | 10 |
| O | 86 | 34.4 |
| Total | 250 | 100 |

**Figure 2.**
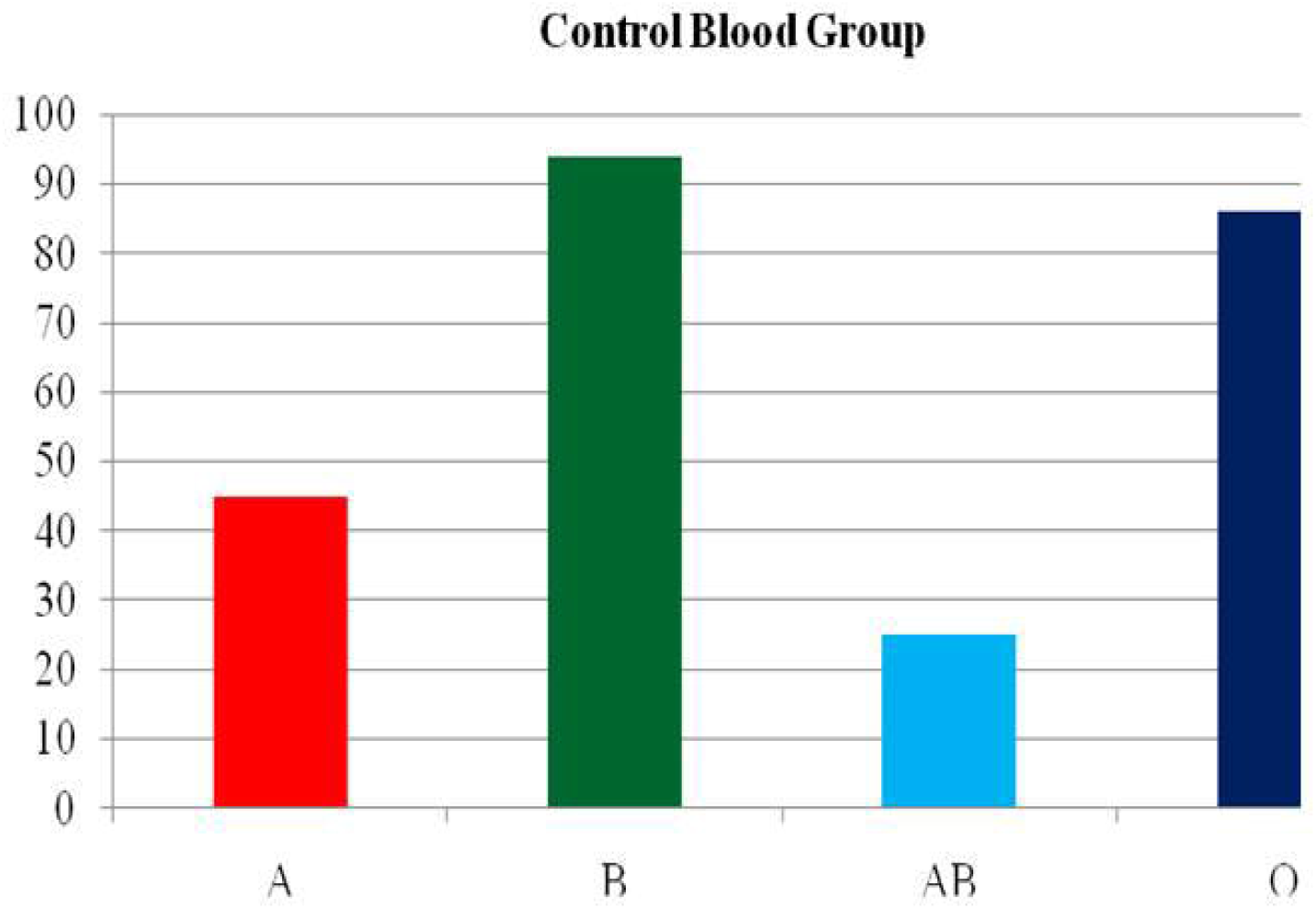
Distribution of ABO blood group in normal population (Control n=250).

**Table 4.** Distribution of ABO blood group in breast cancer patients.

| Blood Group | Number | Percentage | P-Value |
| --- | --- | --- | --- |
| A | 28 | 49.12 | 0.0005 |
| B | 9 | 15.78 | 0.015 |
| AB | 5 | 8.77 | 0.89 |
| O | 15 | 26.31 | 0.382 |
| Total | 57 | 100 |  |

**Figure 3.**
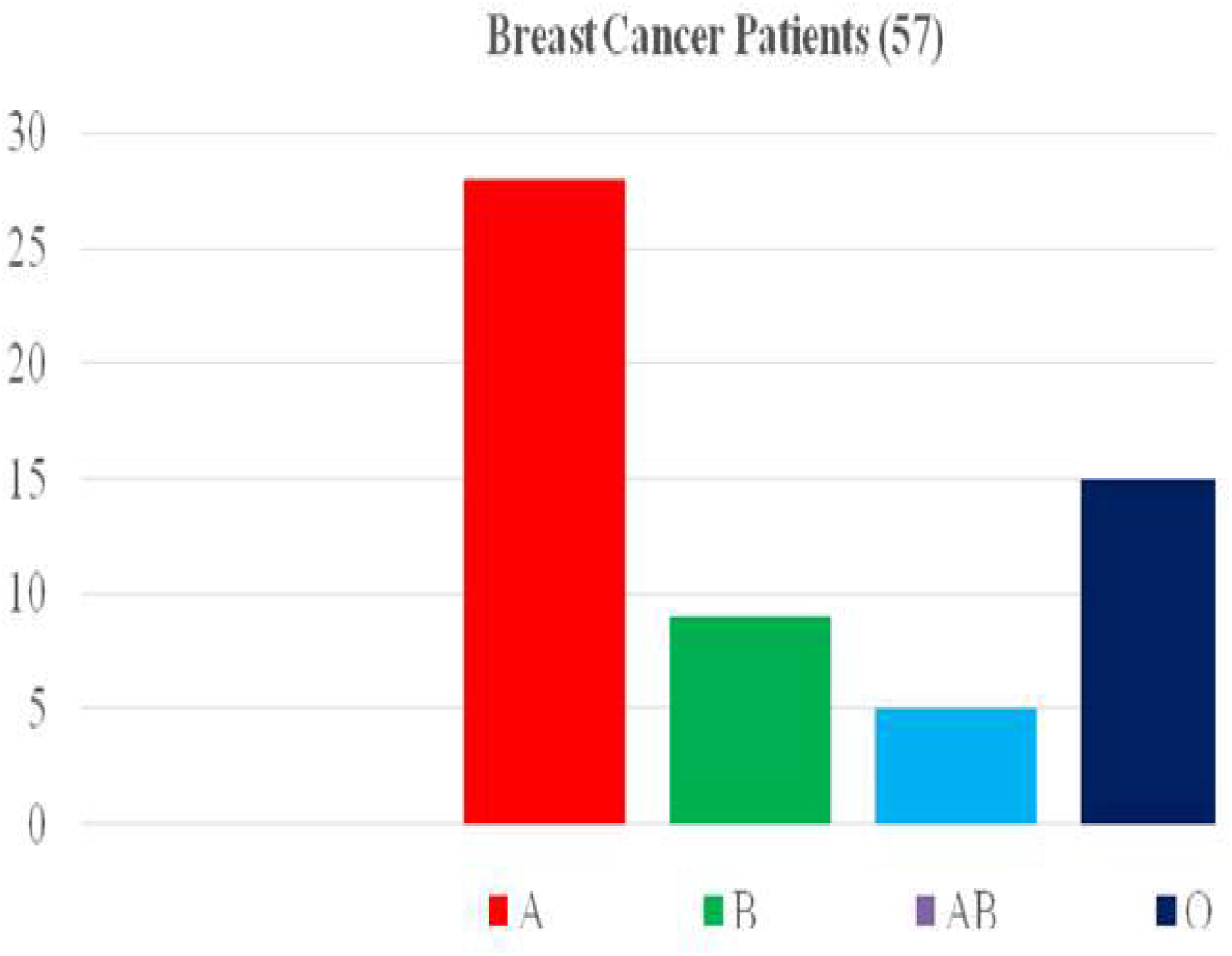
Distribution of ABO blood group in breast cancer patients.

## Discussion

This study aimed to investigate the association between ABO blood groups and risk of cancer. In the present study, data from 243 number of cases and 250 controls, we found significant associations of blood group A with increased risk of Cancer in Eastern (U.P.). Studies across the word has shown association of blood group A with, Breast Cancer (18–21), pancreatic cancer (22–25), liver Cancer (26) gastric and lung Cancer (27,28,29,32,33,34) risk. Data from a these studies have shown that blood group A is associated with breast cancer, liver and pancreatic cancer, and lung cancer risk.

## Conclusion

In conclusion, is found that different ABO blood groups are associated with different type of diseases. Our study also showed that blood type A was more associated to cancer patients and blood type AB having least association and risk of cancer patients.

